# Genome-Wide Identification and Comprehensive Analysis of the SBT Gene Family in Soybean (Glycine max)

**DOI:** 10.1101/2024.03.27.586907

**Authors:** Xuan He, Shanhong He, Jiaxin Yang, Zhiyong Yue

**Affiliations:** School of Engineering, Xi’an International University, 18 Yudou Road, Yanta District, Xi’an Shaanxi, 710077, China; School of Medicine, Xi’an International University, 18 Yudou Road, Yanta District, Xi’an Shaanxi, 710077, China

**Author notes:** Corresponding Author:Zhiyong Yue, School of Medicine, Xi’an International University, 18 Yudou Road, Yanta District, Xi’an Shaanxi, 710077, China. These authors contributed equally to this work (Xuan He, Shanhong He).

## Abstract

**Background:** The SBT (Subtilisin-like serine protease) protein, acting as a crucial serine protease, exerts multifaceted roles in orchestrating plant growth, development, and defense responses by modulating cell wall characteristics and activities of extracellular signaling molecules. Despite extensive investigations into SBT family genes in model dicotyledonous plants like Arabidopsis, no studies have yet delved into the SBT gene family in soybean.

**Methods:** This study marks the inaugural comprehensive analysis of the SBT gene family in soybean, uncovering 118 GmSBT members categorized into six distinct subgroups. Additionally, we conducted comprehensive analyses covering syntenic relationships, exon-intron structures, motif compositions, and cis-regulatory elements. Our examination of RNA-seq data revealed distinct expression patterns of various GmSBTs across various above-ground tissues.

**Results:** In essence, this research bridges a critical knowledge gap concerning the SBT gene family in soybean, furnishing foundational insights that will serve as a valuable reference for future investigations into soybean developmental functions and integrated data analysis.

## Introduction

Soybean, an essential oilseed crop in the leguminous plant family, is extensively cultivated across the United States, South America, and East Asia. Renowned for its remarkable ability to fix biological nitrogen through symbiotic relationships with soil rhizobacteria, soybean serves as a crucial nitrogen source, supplying vital protein feed for animals and oil for human consumption (Masuda and Goldsmith, 2009).The advent of transcriptome sequencing technology has facilitated the acquisition of complete genetic information of the soybean genome through whole-genome shotgun sequencing methods. Furthermore, RNA-Seq has been instrumental in generating valuable genetic resources, such as a soybean gene atlas, which aids in exploring the genetic underpinnings of soybean biology (Schmutz et al., 2010; Liu et al., 2020).Bioinformatics analyses leveraging soybean genomic data have significantly advanced systematic studies on soybean gene family evolution and molecular characterization.

The precise regulation of protein levels is essential for the normal functioning of plant cells, with protein level regulation hinging on the delicate balance between protein synthesis and degradation. Protein hydrolysis, being essentially irreversible, involves proteinases selectively degrading proteins to modulate various aspects of plant growth, development, and defense mechanisms (Schaller, 2004). The majority of plant proteinases belong to the catalytic serine peptidase type, with Bacillus subtilis-associated proteinases constituting the largest family among them. These proteinases, typically expressed in plants under stress conditions, play a vital role in autophagy processes (Paulus et al., 2020). Consequently, understanding the role of the SBT gene family in regulating plant growth, development, and responses to environmental stimuli holds significant importance.

Subtilisin-like serine proteinase (SBT), the second-largest member of the serine proteinase family, exhibits the capability to hydrolyze proteins into peptides, which serve as signaling molecules or ligands binding to receptors, thereby participating in signal transduction pathways (Havé et al., 2018). SBTs are widely distributed across various organisms, including plants, fungi, bacteria, and parasites. The SBT protein family is characterized by a highly conserved structural domain, peptidase_S8 (PF00082), featuring a conserved catalytic triad comprising aspartic acid (Asp), histidine (His), and serine (Ser), serving as specific substrate binding sites. Additionally, plant SBTs have been found to possess proteinase-associated (PA) domains (PF02225) and Inhibitor_I9 domains (PF05922) (Schaller et al., 2018). The first plant-based Bacillus subtilis proteinase was cloned in melon, followed by systematic analysis of SBT gene families across various species (Yamagata et al., 1994). In Arabidopsis thaliana, for instance, 56 SBT family members have been identified, with specific members such as AtSBT6.1 and AtSBT6.2 regulating cell elongation by processing GOLVEN1 peptides, and AtSBT3.8 being involved in the biosynthesis of biologically active PSK peptides, thereby enhancing osmotic stress tolerance in transgenic plants upon overexpression (Rautengarten et al., 2005; Stührwohldt et al., 2021). Beyond model plants, studies have revealed that overexpression of pineapple AcoSBT1.12 delays flowering time in Arabidopsis under long-day conditions, and knocking down TaSBT1.7 in wheat reduces wheat’s hypersensitive response and resistance to stripe rust (Jin et al., 2021; Yang et al., 2021). These findings underscore the multifaceted roles of SBTs in plant-specific development and responses to environmental stresses, exerting diverse functions in plant growth and defense mechanisms.

Despite possessing a large and complex genome, the identification and analysis of the SBT gene family in the soybean genome remain relatively underreported. This study comprehensively investigates the soybean SBT gene family, encompassing phylogenetic relationships, gene structure, motif composition, promoter elements, chromosomal localization, and protein structure. Subsequent research endeavors further delve into the evolutionary relationships between soybean and other species, as well as expression profiles across different tissues. This study aims to provide an in-depth understanding of the functions of the GmSBT gene family, laying the groundwork for future investigations into the mechanisms of GmSBT genes in soybean development.

## Results

### Identification and Characterization of SBTs in Soybean

Following the amalgamation of search results from HMMER and BLASTP, and subsequent confirmation of the target structural domains using NCBI-CDD, a total of 118 GmSBT genes were pinpointed within the soybean genome. They were designated based on their phylogenetic relationships with Arabidopsis. The length of GmSBT proteins ranged from 118 to 1427 amino acids, corresponding to molecular weights spanning from 1.22 kDa to 152.92 kDa. Their isoelectric points ranged from 4.83 to 9.64, indicating their weakly basic nature. The instability index ranged from 16.8 to 44.27, with the majority of GmSBT proteins exhibiting stability. Furthermore, the aliphatic index ranged from 65.41 to 106.25, signifying significant thermal stability of these proteins. The total average hydrophobicity ranged from −0.398 to 0.353, with most GmSBT proteins displaying hydrophilicity.

Subcellular localization prediction analysis unveiled that GmSBT proteins are predominantly localized in chloroplasts (55), extracellular space (10), endoplasmic reticulum (10), vacuolar membrane (12), plasma membrane (11), cytoplasm (10), cytoskeleton (5), nucleus (3), mitochondrion (1), and peroxisome (1). This suggests that GmSBT proteins may primarily express and function in these organelles. These insights contribute to a comprehensive understanding of the structural and functional attributes of identified GmSBT genes in soybeans (Table 1).

### Phylogenetic Analysis of Soybean SBTs

To investigate the evolutionary relationships of GmSBT genes, we employed the neighbor-joining method to construct a phylogenetic tree comparing GmSBT genes with Arabidopsis SBT genes. Full-length protein sequences of 56 Arabidopsis SBTs (AtSBTs) and 118 soybean SBTs (GmSBTs) were aligned to generate the phylogenetic tree (Figure 1). Subsequently, soybean SBT genes were classified into six subfamilies, designated as Group I to Group VI. Notably, Group I emerged as the largest subfamily, encompassing 9 AtSBTs and 54 GmSBTs. The substantial disparity in SBT gene numbers between Arabidopsis and soybean suggests that Group I SBTs in soybean may harbor a more diverse array of functions, possibly undergoing evolutionary divergence between dicotyledonous and monocotyledonous plants.Interestingly, Group III exhibited a significantly lower number of GmSBTs compared to AtSBTs, suggesting potential contraction within this group.

**Figure 1.**
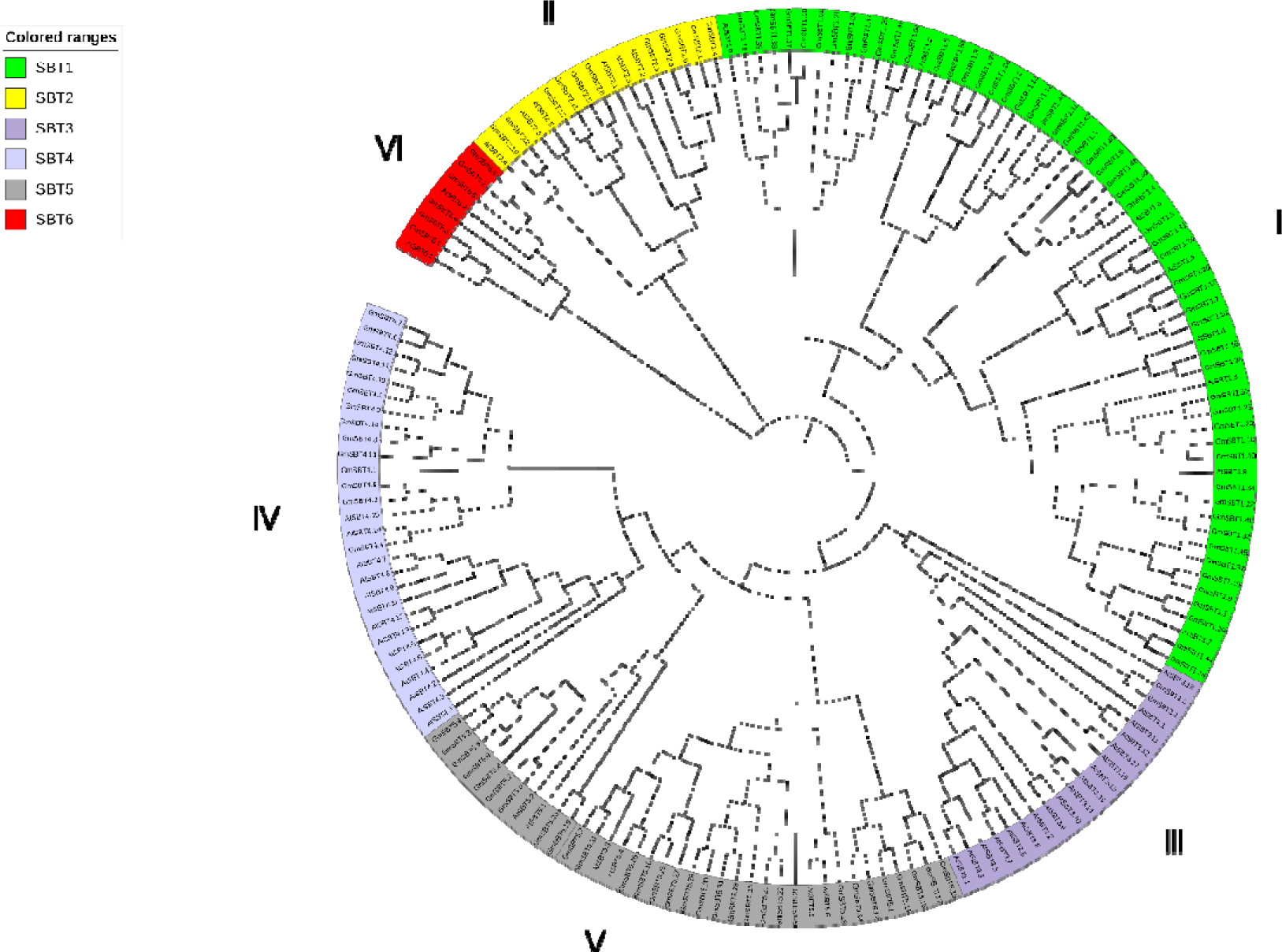
Phylogenetic relationship of the SBT proteins of Gm (Glycine max), At (Arabidopsis thaliana). the evolutionary tree of the GmSBT family was constructed using the neighbor-joining method.

Further exploration of SBT gene evolution between monocots and dicots was conducted based on the phylogenetic tree with maximum likelihood method. A total of 409 SBT genes were categorized into 6 groups, with members of the same class tending to cluster within specific subfamilies. Notably, Group I emerged as the largest subfamily with 199 members, indicating potential divergence among members of this group between monocots and dicots.Conversely, Group VI represented the smallest subfamily with only 14 genes, reinforcing the notion that this subfamily is conserved in evolutionary differentiation (Figure 2). These findings shed light on the evolutionary dynamics of SBT genes between dicots and monocots, offering valuable insights into their functional divergence and conservation across plant species.

**Figure 2.**
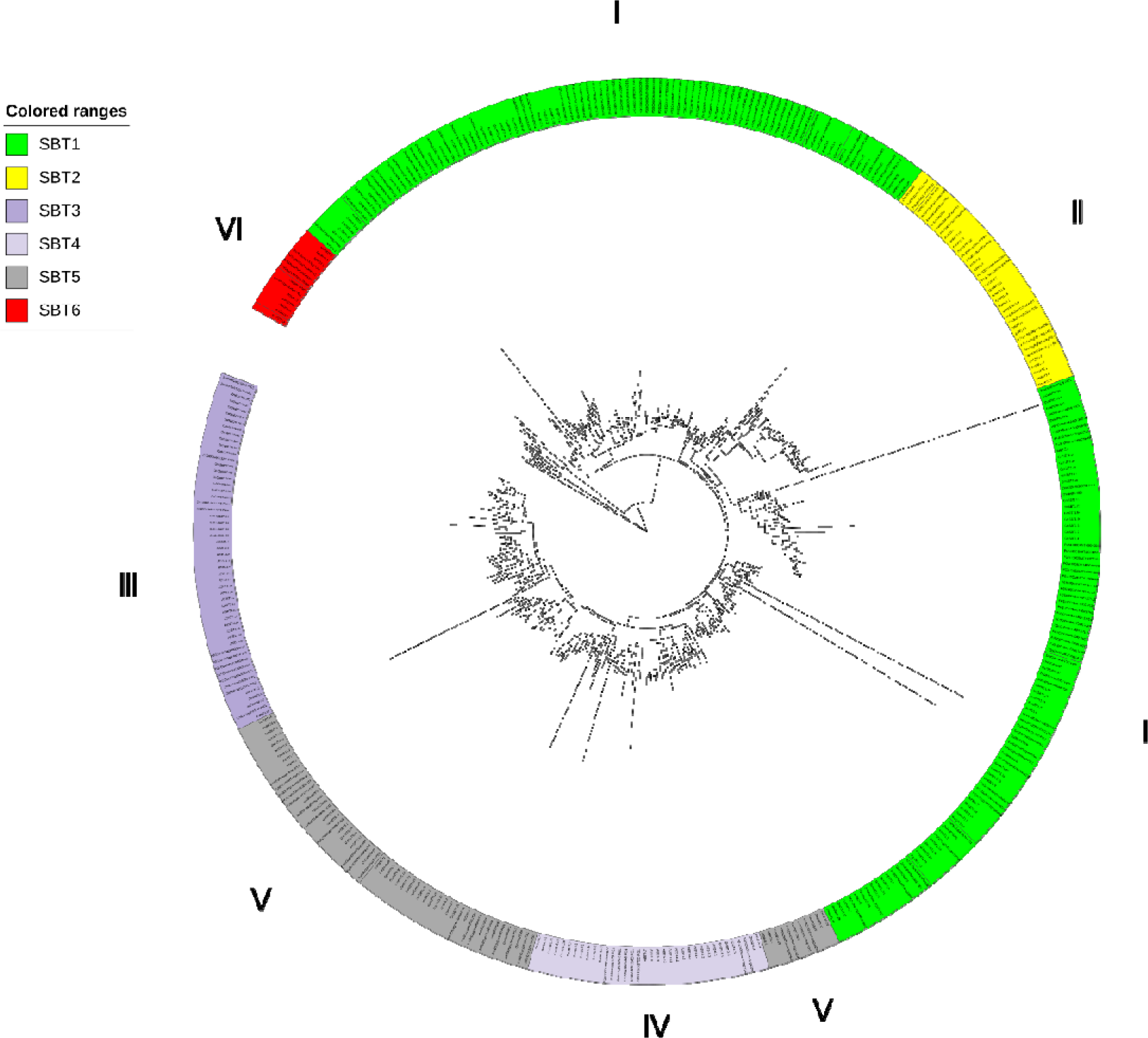
Phylogenetic relationship of the SBT proteins of Gm, At, St(Solanum tuberosum), Os (Oryza sativa), and Zm(Zea_mays).The interspecific evolutionary tree of SBTs was constructed using the maximum likelihood method with 1,000. The phylo-genetic tree divides SBTs into six subfamilies, and the subfamilies are represented with different colors.

### Chromosome Location and Evolutionary Analyses of *GmSBTs*

The chromosomal localization analysis of GmSBT revealed an uneven distribution of 118 TaSBT genes across 20 chromosomes (Supplementary Figure 1). In order to probe the potential mechanisms driving the amplification of GmSBT, we investigated gene duplication events intrinsic to soybean itself. A total of 103 segmental duplications were identified, indicating the pivotal role played by segmental duplications in the expansion of SBT genes within the soybean genome (Figure 3). Recognizing the significant contribution of comparative syntenic maps to evolutionary studies, we also constructed two soybean comparative syntenic maps pertaining to Arabidopsis thaliana, potato, rice, and maize species (Figure 4). In dicot comparisons, we observed 48 homologous gene pairs shared between soybean and Arabidopsis, and 68 such pairs between soybean and potato. Conversely, in monocot comparisons, soybean exhibited 23 homologous gene pairs with rice and 28 with maize, indicating substantial conservation of these genes throughout evolution. We computed Ka, Ks, and Ka/Ks ratios for these gene pairs (Supplementary Table 1) and noted that the Ka/Ks ratios for all SBT gene pairs were less than 1, providing further evidence that the SBT gene family has been subjected to strong evolutionary selection pressure.

**Figure 3.**
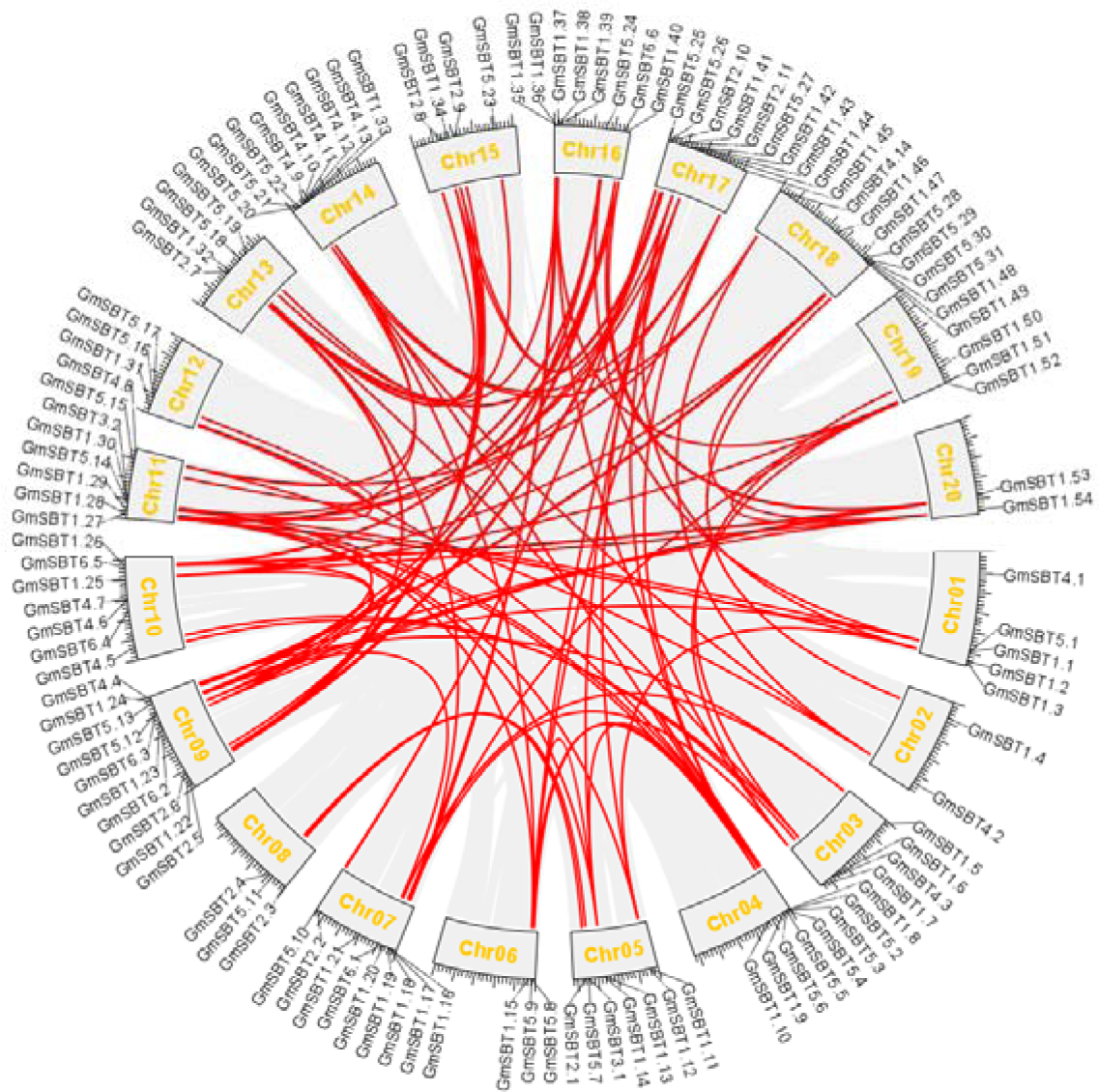
Synteny analysis of SBT genes in soybean. Schematic representation for both chromosomal distribution and interchromosomal relationships of SBT genes.

**Figure 4.**
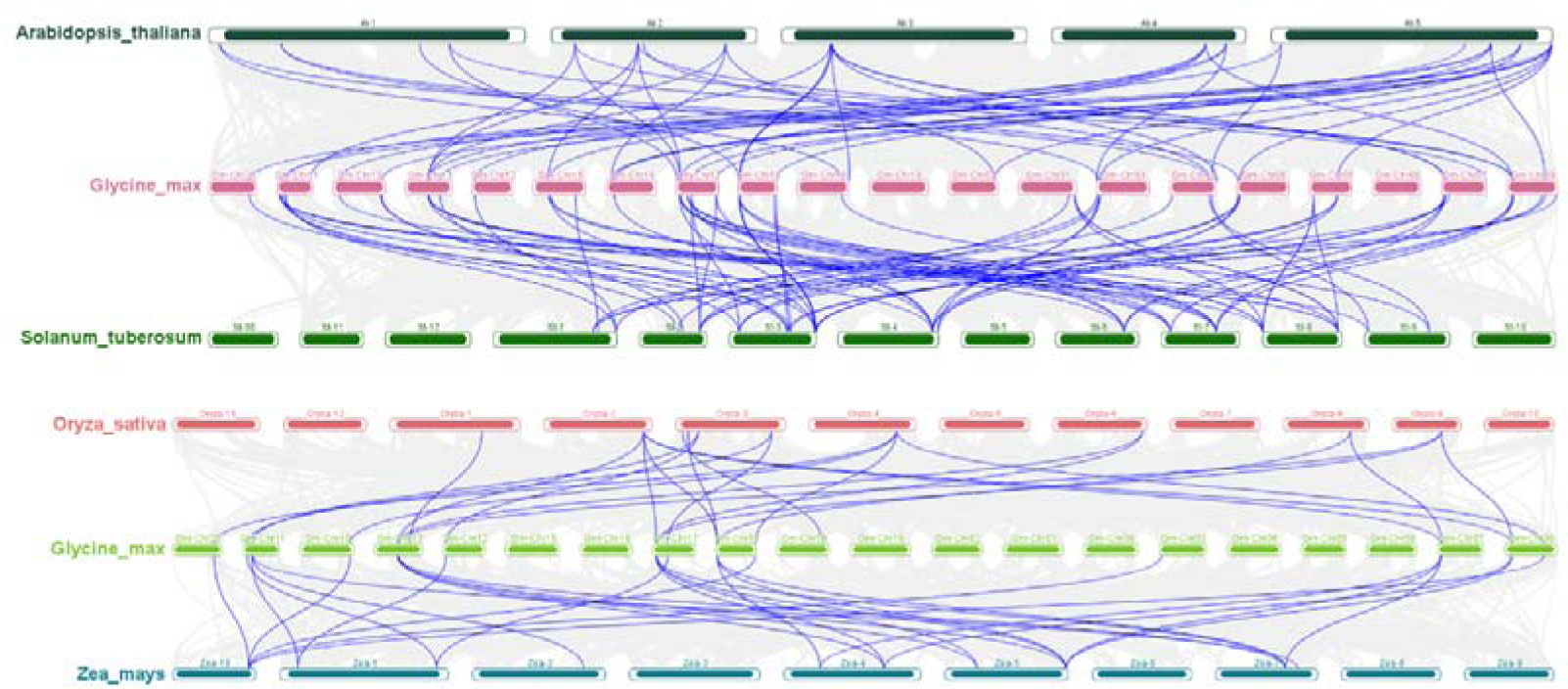
Synteny analysis of SBT genes between soybean and Arabidopsis, potato, rice, and maize. (A) Synteny analysis of SBT genes between soybean and two dicotyledonous plants, Arabidopsis and potato. (B) Synteny analysis of SBT genes between soybean and two monocotyledonous plants, rice and maize. Gray lines in the background indicate the collinear blocks within soybean and other plant genomes, while the blue lines highlight the syntenic SBT gene pairs.

**Figure 5.**
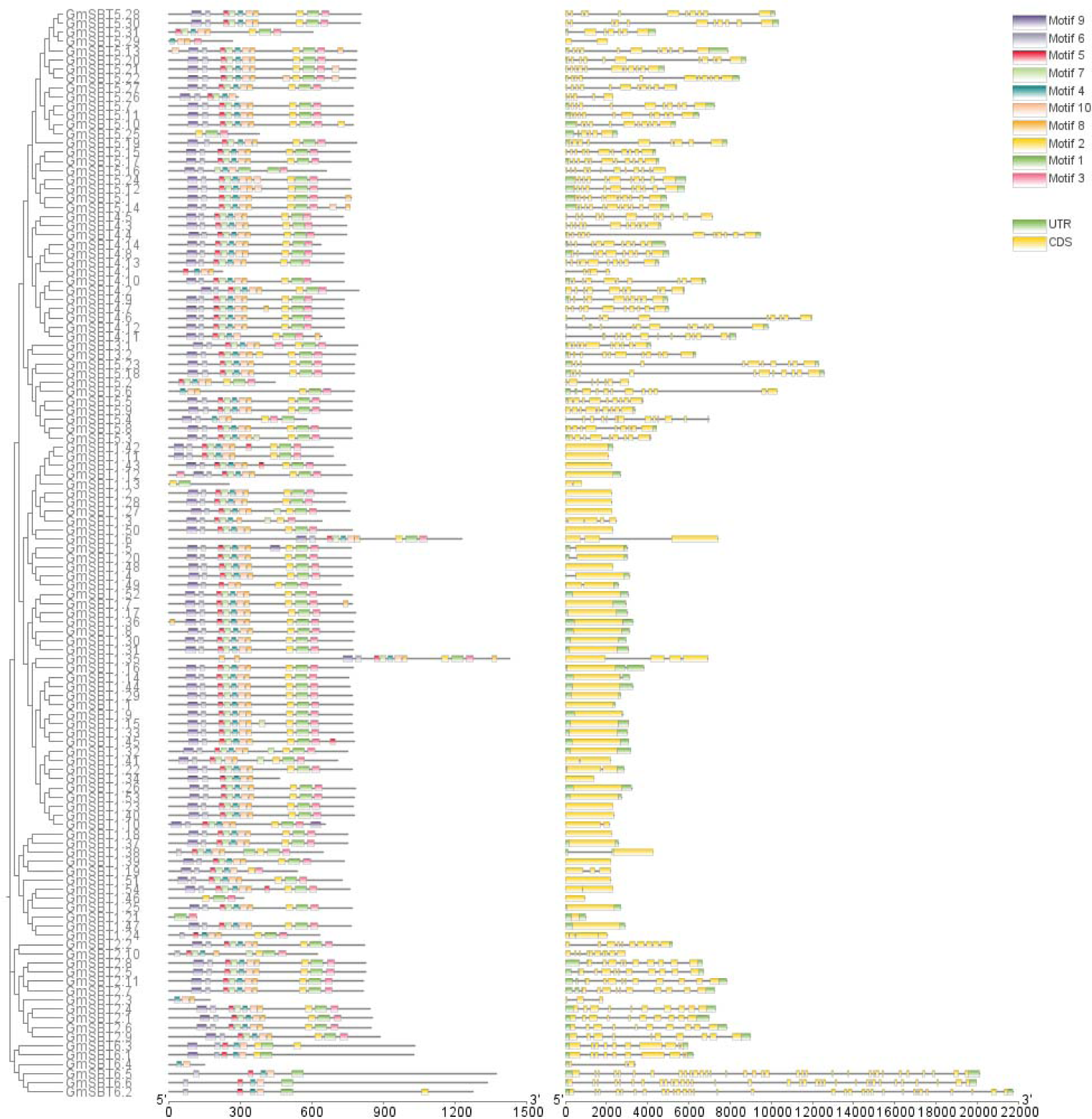
Genetic structure of the SBT gene family in Glycine max. (A) Phylogenetic tree of the SBT gene family. (B) Motif pattern of SBT gene family. (C) SBT gene family exon structure diagram.

### Gene Structure Characterization and Protein Motif Identification

To delve deeper into the structural variances and evolutionary connections among GmSBT genes, we utilized the Neighbor-Joining (NJ) method to construct the GmSBT phylogenetic tree. We then conducted an analysis of the gene structure, conserved domains, and motifs of GmSBT genes (Figure 6A). Motif structures within protein molecules play specific functional roles, and the identification of these motifs can elucidate the characteristics of the gene family. It’s noteworthy that most GmSBT genes exhibit 10 motifs, underscoring their conservation (Figure 6B). Evolutionary trees reveal that GmSBT genes sharing similar domain features cluster together in proximity within the GmSBT phylogenetic tree. Furthermore, gene structures exhibit conservation and simplicity, with similar patterns observed within the same group. Coordinated phylogenetic tree analysis indicates that the number and distribution of motifs among members of the same subfamily are consistent, while significant differences in motif distribution exist among gene sequences from distinct subfamilies, implying potential divergences in gene function (Figure 6C).

**Figure 6.**
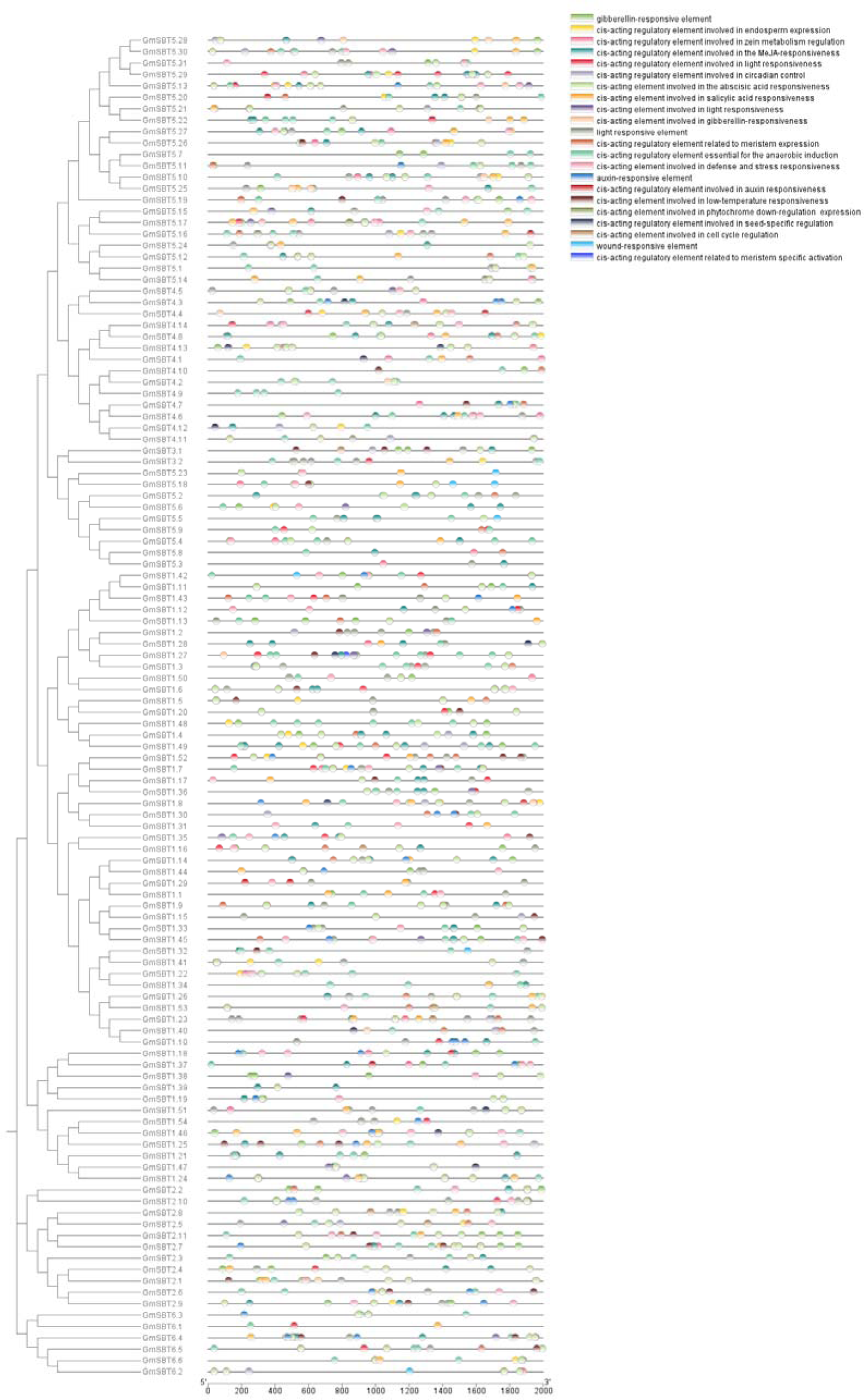
Putative regulatory cis-elements in soybean SBT family gene promoters.

### *Cis*-Acting Elements Identification in Promoter Region of GmSBTs

The 2000 bp upstream promoter sequence of the GmSBT gene’s start site was examined, utilizing the PlantCARE online database to analyze its cis-acting elements, including multiple CAAT boxes and TATA boxes. This analysis confirmed their ability to facilitate normal transcription. Our investigation revealed that the cis-acting regulatory elements within the promoter regions of GmSBT genes primarily fell into four categories: light-responsive, hormone-responsive, stress-responsive, and growth and metabolic-responsive elements (Figure 7). Notably, stress-responsive cis-elements within these promoters played roles in drought response, low-temperature adaptation, defense mechanisms, and anaerobic conditions. Furthermore, we identified cis-elements associated with salicylic acid (SA), methyl jasmonate (MeJA), gibberellins (GA), and auxin (IAA).

**Figure 7.**
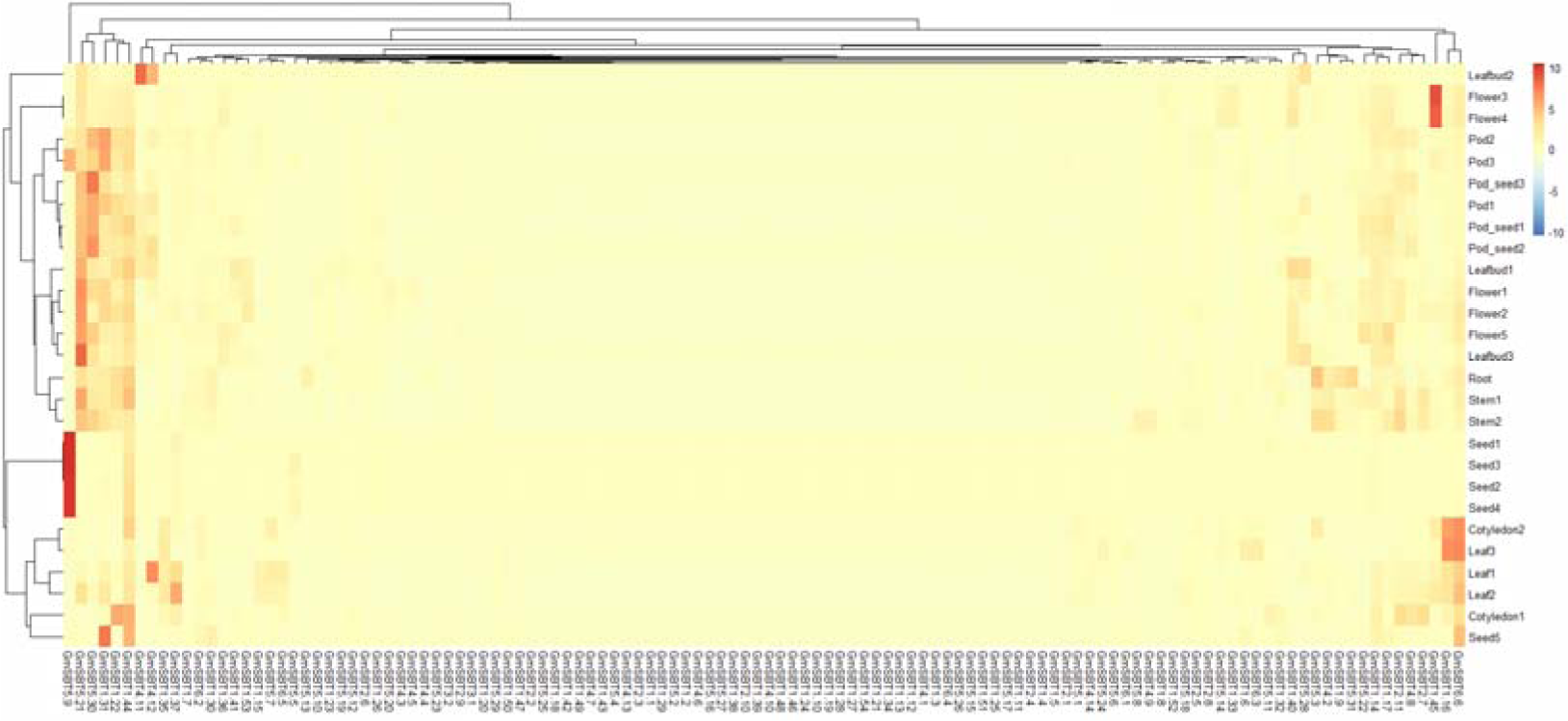
Expression pattern analysis of the soybean SBT family genes in different tissues and organs using transcript data. Expression patterns of the 118 genes were investigated in 27 different tissues and organs using publicly transcriptome data.

### Expression profiles of *GmSBT* genes at different developmental stages

To explore the potential role of GmSBT genes in soybean growth and development, we conducted an analysis of the expression profiles of all SBT genes across various tissues and developmental stages using publicly available transcriptome databases. Our investigation revealed that the 118 GmSBTs display diverse organ-specific expression patterns, offering insights into the functional divergence of the SBT gene family in soybean growth and development (Figure 8, Supplementary Table 2). Notably, genes belonging to Group I, Group V, and Group VI exhibit distinct expression profiles across different tissues, suggesting their involvement in soybean development. Conversely, Group III genes were found to be consistently poorly expressed across all tested samples, indicating a potential lack of involvement in soybean developmental processes. Furthermore, five genes (GmSBT5.9, GmSBT5.21, GmSBT5.30, GmSBT1.22, GmSBT1.44) demonstrated ubiquitous expression across all tissues analyzed, implying essential roles in both vegetative and reproductive growth in soybean.

**Figure 8.**
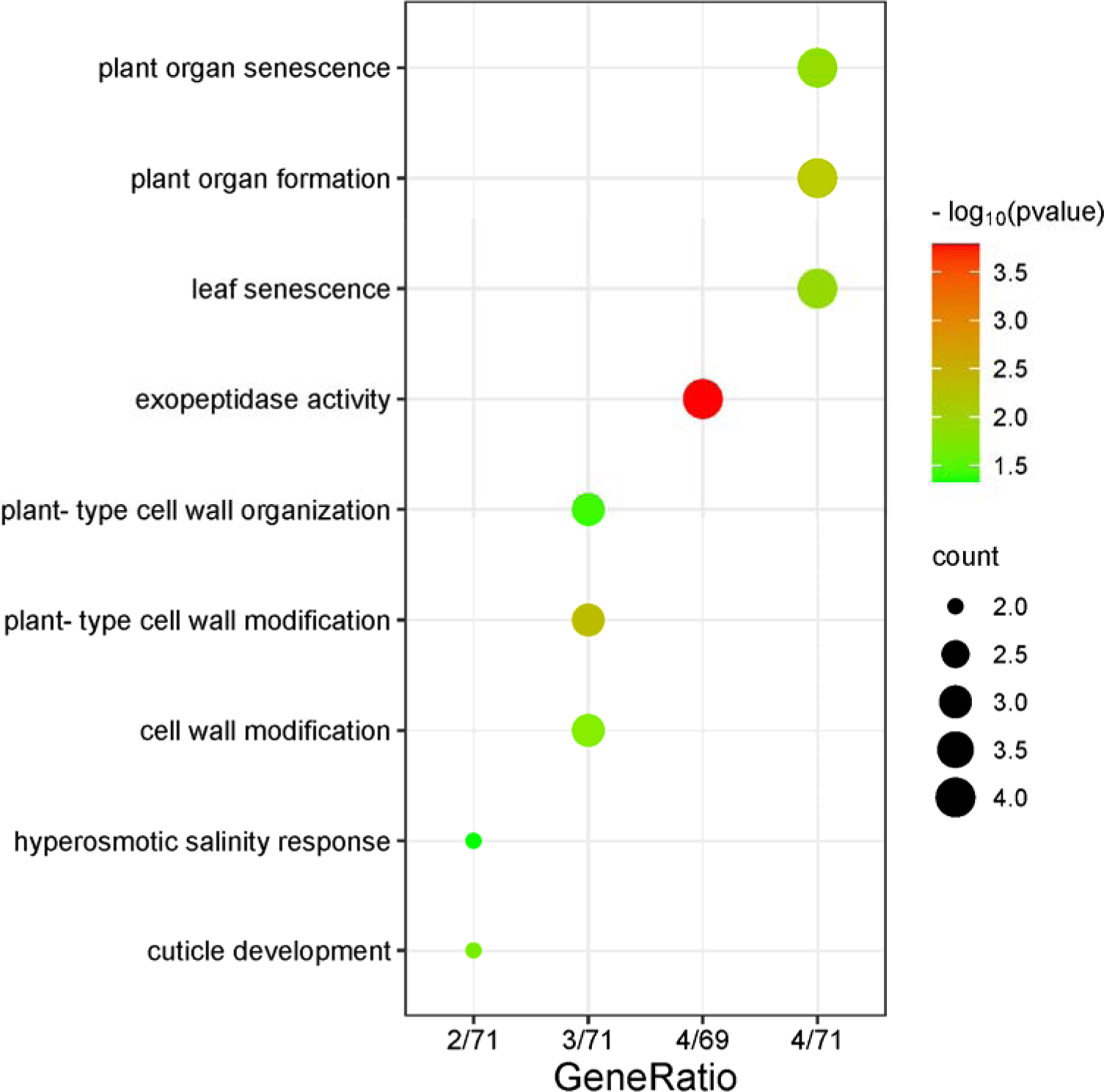
GO enrichment analysis of GmSBTs. Each row corresponds to a valid GO item; The list shows GeneRatio (genes enriched in differentially expressed genes in the pathway/functional genes in differentially expressed genes). The bubble size represents the number of genes, and the color gradient represents—log10 (p-value).

### GO analysis

To unravel the regulatory pathways of these GmSBTs, we conducted GO enrichment analysis. This analysis revealed 63 distinct GO terms spanning biological processes (BP), molecular functions (MF), and cellular components (CC) (Supplementary Table 3, Figure 9). Notably, we observed that GmSBTs are primarily associated with the formation processes of plant organs, including plant-type cell wall tissue, plant organ senescence, and plant organ formation. Additionally, GmSBTs are implicated in the regulatory processes of serine peptidases, exemplified by serine-type endopeptidase activity, serine-type peptidase activity, and endopeptidase activity.

## Discussion

The subtilisin-like serine protease from Bacillus subtilis is a protein digestive enzyme originally obtained from Bacillus subtilis spores. The mature form is a spherical protein containing 275 residues, with multiple α-helices and a large β-folded sheet. Most proteases in plants belong to the catalytic serine peptidase type. Within the serine peptidases, those associated with the bacterial subtilisin-like serine protease constitute the largest family, thus, studying the role of the SBT gene family in regulating plant growth and development, responding to environmental regulation, and addressing biological stress is crucial (Schaller et al., 2018). It has been reported to be involved in various cellular processes, such as protein activation in many plants. This unique region is used as a binding site for specific substrates (Schaller et al., 2012).

The SBT gene family exists throughout the plant kingdom and plays various roles in plant growth and defense. The SBT family has been extensively studied in Arabidopsis thaliana, with a total of 56 AtSBT family members identified. SBT proteins have been reported to be widely present in plants, fungi, bacteria, parasites, etc., and are relatively conserved across different plant species. Genome-wide analysis of the SBT gene family has been performed in *Arabidopsis thaliana*, *Vitis vinifera*, pineapple, *Zea mays,,Solanum tuberosum* And *Musa accuminta* (Rautengarten et al., 2005; Cao et al., 2014; Norero et al., 2016; Wang et al., 2023; Purwar et al., 2024).In this study, 118 GmSBT genes were identified from the soybean reference genome using the HMM model and blastp alignment. The proteins encoded by these genes contain 118-1427 amino acids, with molecular weights ranging from 1.22 to 152.92 kDa and isoelectric points ranging from 4.83 to 9.64, indicating that these proteins are weakly basic. The instability index ranges from 16.8 to 44.97, with most GmSBT proteins showing stability. The aliphatic index ranges from 65.41 to 106.25, indicating significant thermal stability of these proteins. The grand average of hydropathicity ranges from −0.398 to 0.353, with most GmSBT proteins exhibiting hydrophilicity. Proteins encoded by SBT family members have different physicochemical properties, functions, and regulatory mechanisms, but they all share a stable SBT domain. Subcellular localization results indicate that most of these proteins are localized in chloroplasts. This information indicates that members of the GhSBT family likely play pivotal roles in numerous biological processes, encompassing plant growth and development.

To study the phylogenetic relationship of soybean SBT genes, a phylogenetic tree was constructed. According to the phylogenetic relationship, GmSBT members were divided into 6 groups. Studies on gene duplication events illustrate the potential dispersal mechanism of GmSBT, indicating that duplicated genes often come from the same subfamily. Gene structure and motif composition further demonstrate the relative conservation among members of the same subfamily, consistent with previous studies. Additionally, collinearity analysis reveals a large number of tandem repeats between homologous chromosomes. These results indicate that the pattern of gene family expansion, including replication and multiple gene copies, can prevent loss of gene function caused by gene mutations, highlighting the importance of their function (Cannon et al., 2004; Xu et al., 2012). We calculated the Ka/Ks ratio of gene pairs, and the Ka/Ks ratio of all SBT gene pairs is less than 1, further confirming that the SBT gene family has undergone strong evolutionary selection pressure, which is of great significance for understanding the evolution of soybeans (Panchy et al., 2016).

During exposure to abiotic stresses, the cis-acting elements located upstream of each gene assume a significant role. Analysis of these cis-acting elements revealed that GmSBTs potentially participate in various responses(William Roy and Gilbert, 2006). including light responsiveness, stress responsiveness, growth and metabolic responsiveness, as well as hormone responses such as those to abscisic acid, gibberellin, salicylic acid, growth hormone, and MeJA. These findings suggest that GmSBTs are not only engaged in multiple signaling pathways but also contribute to plant growth, development, and defense responses(Schaller et al., 2018). This provides a valuable reference for screening stress resistance genes. Validation of the spatiotemporal expression profiles from plant tissue data revealed that the expression pattern of the GmSBT gene exhibited a distinct organ-specific tendency, with certain subgroups displaying low expression levels, consistent with prior research findings(Beers et al., 2004). In conclusion, the valuable insights garnered from this study will enhance our understanding of soybean SBT growth and development. Future research endeavors should delve deeper into elucidating the molecular mechanisms underlying soybean SBT during growth and development.

### Conclusion

In the present study, a genome-wide identification of the soybean SBT gene family led to the initial characterization of 118 GmSBT members. Extensive bioinformatics analyses laid the groundwork for further exploration of this gene family. Expression profiling indicated that the gene family plays a significant role in controlling plant growth and development, and this research provided a foundation for understanding the biological functions of SBT genes in soybean. In summary, these findings offer valuable information for advancing investigations into the roles of GmSBTs in plant development and growth.

## Materials and Methods

### Identifcation and Characterization of *SBT* genes in soybean

The Hidden Markov Model (HMM) (Potter et al., 2018) for the Peptidase_S8 domain (PF00082) (Wen et al., 2020) was acquired from the Pfam database (Bateman, 2004) and utilized as a query to interrogate the soybean genome database retrieved from Ensembl Plants(Bolser et al., 2015). Concurrently, SBT protein sequences from Arabidopsis thaliana were sourced from the Arabidopsis Information Resource (TAIR) database (Lamesch et al., 2012) and subjected to BLASTP comparisons against the soybean protein database. Following the integration of HMMER and BLASTP search outcomes, each chosen candidate underwent further scrutiny for the presence of the conserved domain using the NCBI Conserved Domain Database (CDD) (Marchler-Bauer et al., 2011). To ascertain additional protein properties, ExPASy tools were employed for calculating the molecular weight (MW) and isoelectric point (pI) of the GmSBT proteins(Gasteiger et al., 2005), while WoLFPSORT facilitated subcellular localization predictions(Horton et al., 2007).

### Phylogenetic analysis of soybean SBT

All identified soybean SBT protein sequences were subjected to multiple sequence alignments with Arabidopsis SBT sequences collected from TAIR, employing Muscle with default parameters. The Ensembl Plants database was used to obtain the whole genome and protein sequences of potato(*Solanum tuberosum*), rice (*Oryza sativa*), and maize (*Zea mays*) using the HMM-SBT model. The phylogenetic analysis was conducted using MEGA11 (Dejosez et al., 2023), adopting the neighbor-joining statistical method and maximum likelihood under default settings (except for bootstrapping with n = 1,000 replicates).Based on the clustering scheme of GmSBT, a total of 118 GmSBT sequences were classified into sex distinct groups. The evolutionary tree was visualized using the online tool iTOL (Letunic and Bork, 2021).

### Chromosome Distribution and Evolutionary Analyses of soybean SBT genes

From the soybean genome annotation files, we extracted the chromosomal locations of GmSBT genes. Subsequently, we conducted all-against-all comparisons within the soybean genome using BLASTP searches with an E-value threshold less than 10^(−10) to elucidate information about gene pairs in a syntenic analysis. Based on the BLASTP results, we analyzed syntenic blocks and gene duplication events using McScanX within the TBtools software suite (Chen et al., 2023). To visually represent the relationships among GmSBT gene pairs, an advanced circular plot was generated using the Advanced Circos plugin within Tbtools (Chen et al., 2022), with duplicated GmSBT gene pairs depicted by interconnected solid black lines. Furthermore, we calculated Ka and Ks values for the duplicated GmSBT gene pairs using the Ka/Ks calculator embedded in TBtools to assess the divergence among the duplicated genes (Chen et al., 2023).

### Gene Structure, Motif Analysis, and *Cis*-Acting Elements Identification

Utilizing the TBtools program, we depicted the gene structure features and exon-intron organization of GmSBT based on the full-length genomic sequences and protein-coding sequences of the provided genes (Chen et al., 2023). To identify potential motifs, we employed the MEME (Multiple Expectation Maximization for Motif Elicitation) web-based motif discovery server (Bailey et al., 2015), setting parameters including motif width < 50, maximum number of motifs < 20, and e-value cutoff < e^(−5. For each GmSBT, we selected a 2.0 kilobase upstream sequence of the gene as the promoter region, which was then extracted from the soybean genome. Subsequently, we utilized the PlantCARE website to predict cis-regulatory elements within these promoter regions(Lescot et al., 2002).

### Expression pattern of *SBT* genes in various tissues

To analyze the expression patterns of GmSBT genes under diverse conditions, encompassing various growth and developmental stages across multiple plant tissues, we obtained RNA-seq data from the SoyMD databases (Yang et al., 2023). The annotated information furnished by the Glycine max Reference Genome served as the reference database for analyzing Gene Ontology (GO). Images were generated using the SRplot website (Tang et al., 2023).

