## Supplementary figures and images for "Genome-Wide Identification and Comprehensive Analysis of the SBT Gene Family in Soybean (Glycine max)"

### Supplemental Figure 1

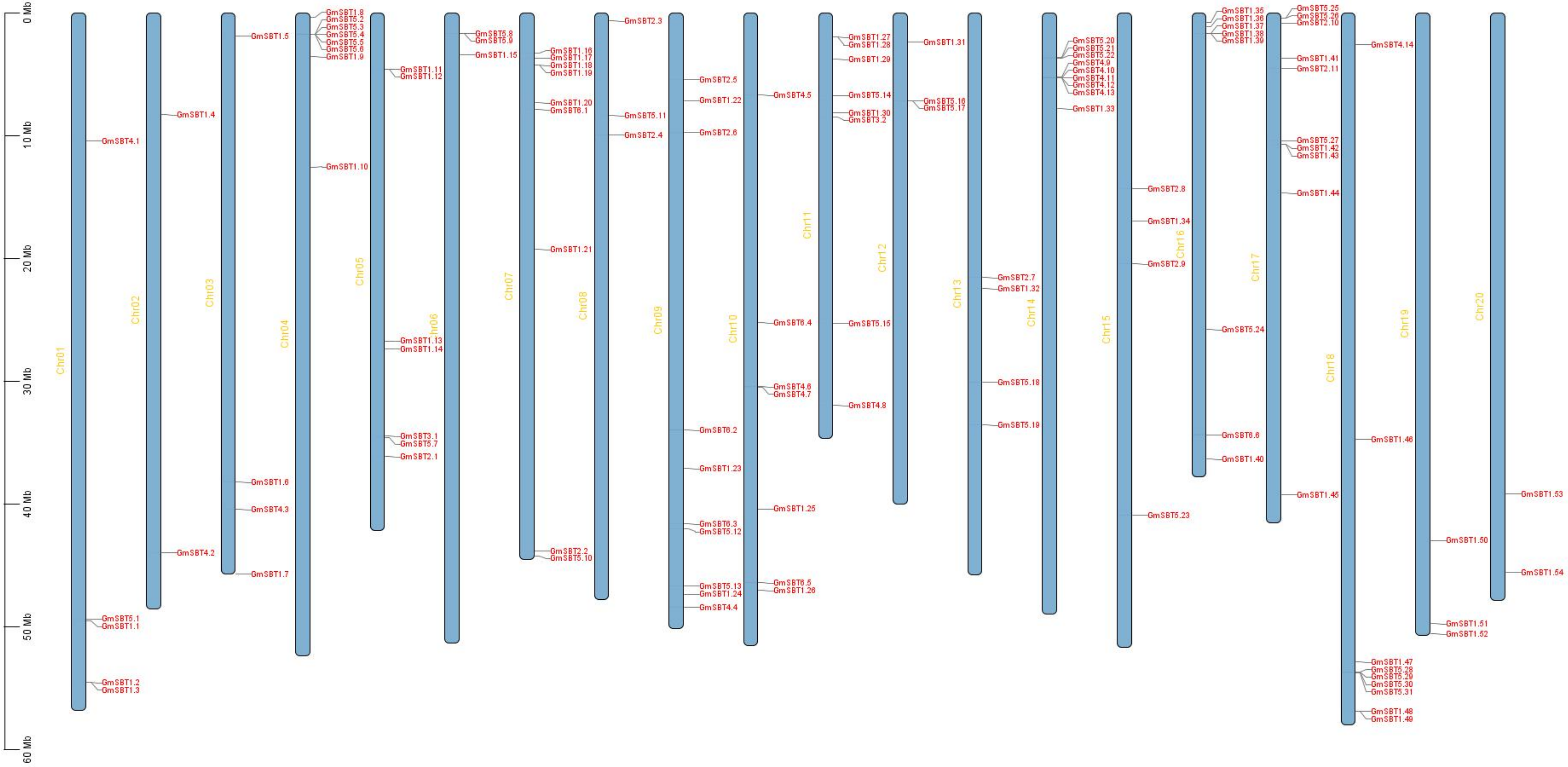
